# PuMA: a papillomavirus genome annotation tool

**DOI:** 10.1101/736991

**Authors:** J. Pace, K. Youens-Clark, C. Freeman, B. Hurwitz, K. Van Doorslaer

## Abstract

High-throughput sequencing technologies provide unprecedented power to identify novel viruses from a wide variety of (environmental) samples. The field of ‘viral metagenomics’ has dramatically expanded our understanding of viral diversity. Viral metagenomic approaches imply that many novel viruses will not be described by researchers who are experts on the genomic organization of that virus. There is a need to develop analytical approaches to reconstruct, annotate, and classify viral genomes. We have developed the papillomavirus annotation tool (PuMA) to provide researchers with a convenient and reproducible method to annotate novel papillomaviruses. PuMA provides an accessible method for automated papillomavirus genome annotation. PuMA currently has a 98% accuracy when benchmarked against the 481 reference genomes in the papillomavirus episteme (PaVE). Finally, PuMA was used to annotate 168 newly isolated papillomaviruses, and successfully annotated 1424 viral features. To demonstrate its general applicability, we developed a version of PuMA that can annotate polyomaviruses.

PuMA is available on GitHub (https://github.com/KVD-lab/puma) and through the iMicrobe online environment (https://www.imicrobe.us/#/apps/puma)

## INTRODUCTION

Viruses and viral diseases have fascinated humans for millennia (reviewed in (Woolhouse *et al.*, 2012). We are currently in the era of *‘Neovirology’* (Enquist and Editors of the Journal of Virology, 2009), and recent technological advances provide an agnostic approach to sequencing nucleotides from environmental samples. These approaches dramatically accelerated viral discovery (Hurwitz *et al.*, 2016; Nooij *et al.*, 2018). Excitingly, these “viromic” approaches facilitate the identification of highly diverse viruses from a wide array of sample types. However, for these viromic studies to maximize their potential scientific impact, the identified viral genomes should be annotated and curated prior to publication. Many viruses are being identified by researchers who are not necessarily experts on that specific family of viruses thus complicating a detailed curation process. Bioinformatic software packages have the potential to provide quick, accurate, and reproducible annotations. These software tools should be universally available, actively maintained, powerful yet intuitive, and automatable (Rose *et al.*, 2016).

The *Papillomaviridae* (PVs) is a family of viruses with a circular, double-stranded DNA genome about 8,000 bp in length. Prototypical papillomavirus genomes contain six distinct open reading frames (ORFs). The early proteins (E1, E2, E6, E7) manipulate the cellular environment to allow for viral replication. The late L1 and L2 protein are the viral structural proteins. In addition, some viruses contain additional ORFs (e.g., E5, E8, E9, E10)(Van Doorslaer, 2013).The viral upstream regulatory region (URR), is located downstream of the L1 ORF and contains binding sites for viral and host proteins (Bernard, 2013).

We describe a *Papillomaviridae*-specific annotation tool (PuMA). PuMA can be added to computational pipelines, or a graphical user interface is available through the iMicrobe online environment (Youens-Clark *et al.*, 2019). To demonstrate its potential broad usability, we adapted PuMA to annotate viruses belonging to a different taxonomic family.

## METHODS

The papillomavirus annotation tool (PuMA) leverages a set of existing software tools and packages (supplementary table 1). Since PuMA is based on homology-based search approaches, the computer science idiom “garbage in/garbage out” applies. Therefore, PuMA uses the manually curated sequences available in the PaVE database (Van Doorslaer *et al.*, 2013, 2017).

### Annotation of viral proteins

As input, PuMA only requires a viral genome in FASTA format. For consistency, the first nucleotide following the L1 stop codon (i.e. the beginning of the URR) is used as the start of the (circular) viral genome. All open reading frames -a stretch of at least 75 nucleotides located between two stop codons- on the forward strand are translated into putative peptides. A BLASTp homology (Altschul *et al.*, 1990) search against custom databases (available on the PuMA GitHub page) is used to identify the primary viral open reading frames. If available, additional information is used to further improve this process. For example, the consensus start codon of the L1 ORF is located just downstream of a consensus splice acceptor site (Buck *et al.*, 2013). Furthermore, the N-terminus of the majority of L1 proteins contains a conserved MxxWx_7_YLPP motif (x can be any amino acid). Likewise, a multiple sequence alignment step is used to identify the most parsimonious E6 start codon (Fig 1A).

**Figure 1.**
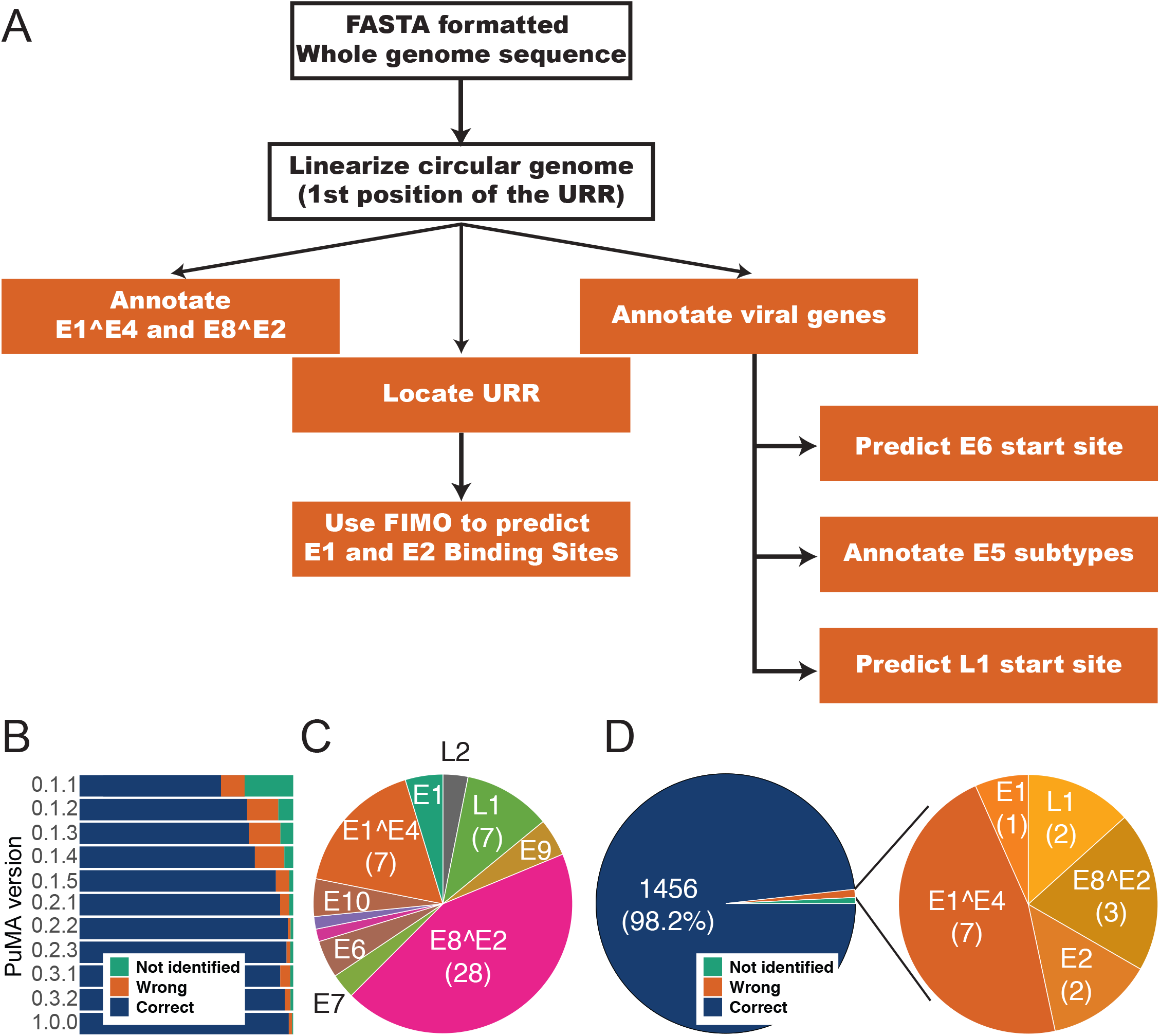
(A) Flowchart illustrating the PuMA algorithm: see methods section for more details. (B) PuMA correctly annotates 98% of manual annotations present on the PaVE database. PuMA annotations were compared to the manually curated genomes in the PaVE database. Iterative changes to the PuMA algorithm improved the accuracy to 98%, however, in the absence of experimental data, PuMA can only provide testable hypotheses. (C) Proteins that were wrongly identified by PuMA v1.0 are plotted. The algorithm primarily miss-identifies spliced proteins (E8^E2 and E1^E4). (D) Figure 4 PuMA correctly annotates novel genomes. PuMA was trained on 481 genomes present on PaVE prior to May 2019. PuMA v1.0 was used to annotate 168 novel viral genomes. These included viruses isolated from human (n=143) and non-human (n=25) hosts. The PuMA annotations were compared to manually curated genome annotations. PuMA correctly identified 98.2% of the viral features. Proteins that were wrongly identified by PuMA v1.0 are plotted.

### Annotation of spliced viral mRNA

A typical papillomavirus transcribes twenty alternatively spliced mRNAs (Graham and Faizo, 2017). At this time, PuMA annotates the two best studied alternatively spliced viral mRNAs, E1^E4 and E8^E2. The splice acceptor is shared between the E1^E4 and E8^E2 mRNAs and is embedded within the viral E2 ORF. We use MUSCLE (Edgar, 2004) to align the newly annotated E2 ORF to its closest previously known relative. The splice acceptor site is identified based on the implied homology between both viruses. A similar homology-based approach is used to identify both splice donor sites located within the E1 ORF (Fig 1A).

### Annotation of *cis* binding sites

Using PaVE derived URR sequences as a training set, MEME (Bailey *et al.*, 2009) generated a consensus E1 (ATDATTGTTGNYAACAAYHAT; D: A, G, or T; N: any base; Y: C or T; H: A, C, or T) and E2 (ACCGNWWNCGGT; W: A or T) binding site profile. PuMA uses FIMO and these profiles (Grant *et al.*, 2011) to identify putative E1 or E2 binding sites within the viral URR (Fig 1A).

### Output

PuMA generates a log file describing the progress of the analysis, a ‘comma-separated values’ (*.csv) file that contains the individual annotations, a ‘general features format’ (*.gff3) formatted file, a GenBank-formatted file, and a pdf file providing a visual representation of the newly annotated genome. PuMA also provides the sequin files needed to streamline submission to GenBank.

## RESULTS AND DISCUSSION

We developed PuMA to generate consistent and uniformly annotated papillomavirus genomes. To test the performance of PuMA, 481 FASTA-formatted viral genomes (with 4,385 associated annotations; Van Doorslaer *et al.*, 2013) were downloaded from PaVE (accessed on 7/29/2018). Iterative changes to the annotation algorithm improved PuMA’s ability to correctly identify viral features. The current version of PuMA (version 1.0), has an accuracy of ~98% (Figure 1B). In the absence of additional information, PuMA will annotate the longest protein in each open reading frame (starting at the first methionine). Many of the discrepancies between PuMA and the (manually) annotated genomes are due to differences in start codon prediction (Fig. 1C). We are currently developing a homology-based approach to improve the identification of initiation codons. Finally, we used PuMA to annotate 168 newly isolated papillomaviruses (Tirosh *et al.*, 2018; Pastrana *et al.*, 2018), and successfully annotated 1424 novel viral features (Fig 1 D) with high accuracy compared to manual annotation.

This homology-based approach assumes that the manually annotated training data is correct. However, since the majority of E1^E4 or E8^E2 encoding transcripts have not been experimentally validated, it is difficult to confirm the biological accuracy of these annotations. Of note, we believe that PuMA corrected 31 E8^E2 proteins previously misannotated on the PaVE database. These corrections will be incorporated in future releases of the PaVE database.

Since PuMA relies on homology-based searches to known viruses, it is easy to adapt PuMA to annotate different viral families. To demonstrate this, we modified PuMA to annotate polyomavirus genomes. Expertly annotated viral genomes were downloaded from PyVE (http://home.ccr.cancer.gov/lco/PyVE.asp). We updated the primary BLAST database to contain polyomavirus sequences (available on GitHub). Since polyomaviruses are translated bi-directionally, we edited the PuMA code to annotate both the forward and reverse strands of the polyomavirus genomes. Without making additional changes, PuMA correctly annotates 87% of polyomavirus proteins compared to the manually annotated PyVE derived genomes. While this accuracy can be improved, this exercise demonstrates that, given a well-annotated database, PuMA can be used to annotate a wide array of viruses belonging to different families.

## CONCLUSION

The goal of PuMA is to provide users with an automated means of uniform and unbiased papillomavirus genome annotation. With the development of new sequencing technologies, PuMA will aid in the labor-intensive process of annotating the provided genome. Even though PuMA is papillomavirus specific, the open source software is combined with the custom functions, the program can easily be adapted for other viruses by simply changing the background databases. This was demonstrated by adapting PuMA to annotate a different family of viruses.

## ACKNOWLEDGEMENTS

The authors acknowledge the members of the Van Doorslaer lab for testing PuMA and useful comments. This research was supported in part by list TRIF startup funds to KVD.

